# Albumin-coated VS1 nanocrystals enable STARD3 inhibition and potentiate fluoropyrimidine therapy in colorectal cancer

**DOI:** 10.64898/2026.07.07.737012

**Authors:** Isabella Caligiuri, Alessandro Bregalda, Gloria Saorin, Luisa MR Napolitano, Giulio Poli, Simona Kranjc Brezar, Urška Kamenšek, Miriana Di Stefano, Kirti S Sonkar, Juan Luis Pacheco-Garcia, Raghurama Hedge, Salvatore Parisi, Jane Budai, Muhammad Adeel, Carlotta Granchi, Marco De Scordilli, Silvia Onesti, Maja Čemažar, Tiziano Tuccinardi, Vincenzo Canzonieri, Flavio Rizzolio

## Abstract

Poor aqueous solubility remains a major obstacle to the translational development of targeted anticancer compounds. VS1, a first-in-class inhibitor of the cholesterol-transfer protein STARD3, has emerged as a promising chemosensitizing agent in colorectal cancer (CRC), but its clinical applicability is limited by its poor water solubility. Here, we combine structural biology, nanotechnology, and functional pharmacology to establish STARD3 inhibition as a delivery-enabled strategy to potentiate fluoropyrimidine therapy. To define the molecular basis of STARD3 inhibition, we solved the crystal structure of VS1 bound to the STARD3 ligand-binding domain at 2.1 Å resolution, revealing direct occupation of the sterol-binding cavity. Molecular dynamics simulations confirmed a stable binding mode and identified the Ω1 loop as a dynamic gate regulating ligand binding and dissociation. To overcome the formulation barrier of VS1, we engineered carrier-free, albumin-coated nanocrystals through sonication-assisted nanocrystallization followed by surfactant exchange with human serum albumin. The resulting rod-shaped nanocrystals displayed nanometric size, narrow size distribution, sustained release, and improved aqueous dispersibility, increasing the apparent solubility of VS1 by more than 14-fold while preserving its molecular integrity and crystallinity. Biologically, VS1 selectively potentiated 5-fluorouracil (5-FU) in CRC cells, with synergistic effects restricted to 5-FU-sensitive models and associated with enhanced reactive oxygen species accumulation. Albumin-coated formulation retained the chemosensitizing activity of the free compound. In HCT-116 xenografts, combined treatment with albumin-coated VS1 nanocrystals and 5-FU significantly reduced tumor growth, prolonged tumor doubling time, and increased intratumoral necrosis without exacerbating systemic toxicity. Together, these findings establish that albumin-coated nanocrystals can overcome the delivery limitations of an insoluble STARD3 inhibitor and provide a formulation-enabled strategy to enhance fluoropyrimidine therapy in colorectal cancer.

**HIGHLIGHTS:**

- A 2.1 Å crystal structure shows VS1 in the STARD3 sterol-binding cavity
- Albumin-coated nanocrystals make the insoluble STARD3 inhibitor VS1 deliverable
- Nanocrystallization raises VS1 apparent solubility >14-fold with sustained release
- VS1 synergizes with 5-FU selectively in 5-FU-sensitive colorectal cancer lines
- Albumin-nanocrystal VS1 plus 5-FU curbs xenograft growth without added toxicity

## INTRODUCTION

Colorectal cancer (CRC) is among the leading causes of cancer-related mortality worldwide, with nearly two million new cases diagnosed annually (GLOBOCAN 2022). Despite advances in early detection and therapeutic strategies, the prognosis for advanced-stage CRC remains poor, with five-year survival rates for metastatic disease remaining below 20% [1]. Current treatment regimens rely heavily on fluoropyrimidine-based chemotherapy, particularly 5-fluorouracil (5-FU), administered either as monotherapy or in combination with agents such as oxaliplatin and irinotecan [2–4]. Although these approaches have improved clinical outcomes, their efficacy is often limited by intrinsic and acquired resistance, as well as systemic toxicity [5–7]. These limitations highlight the need for therapeutic strategies that can restore or enhance fluoropyrimidine sensitivity while avoiding unnecessary toxicity.

Cancer metabolism has emerged as a key determinant of therapeutic response, and cholesterol homeostasis is increasingly recognized as a contributor to tumor progression, membrane signaling, mitochondrial function, and drug resistance [8–10]. Cholesterol is a key component of lipid rafts, specialized membrane microdomains that regulate receptor signaling, intracellular trafficking and drug transport, and its dysregulation has been linked to oncogenic signaling and chemoresistance [2,11,12]. Beyond extracellular cholesterol uptake and de novo synthesis, intracellular cholesterol trafficking is critical for maintaining lipid distribution between organelles and supporting cancer-associated adaptive responses [13,14].

In this context, the StAR-related lipid transfer protein STARD3 has emerged as a potential regulator of cancer cell metabolism. STARD3 localizes to late endosomes and mediates cholesterol transfer through its START domain [15–17]. Overexpression of STARD3 has been reported in several cancer types and is associated with enhanced proliferation, metastasis, and poor clinical outcomes [18–20]. Moreover, STARD3 has been implicated in mitochondrial cholesterol transport and steroidogenesis, processes that may support tumor growth and survival [21,22]. These findings suggest that STARD3 may represent both a biomarker and a therapeutic target in CRC, although its pharmacological modulation remains largely unexplored.

In previous work, we identified a small-molecule STARD3 inhibitor, termed VS1, through in silico screening [20,23]. VS1 binds to the STARD3 START domain with micromolar affinity and exhibits antiproliferative activity in cancer cell lines. However, its mechanism of action at the structural level and its therapeutic potential in CRC have not been fully characterized. More importantly, its development is hindered by a major pharmaceutical barrier: VS1 is poorly soluble in water, compromising aqueous dispersibility, systemic administration, and meaningful in vivo evaluation. This limitation is common to many potent anticancer chemical molecules identified through target-based discovery campaigns, where pharmacological activity is not matched by suitable formulation properties.

Drug nanocrystal technology offers a direct, carrier-free strategy to address this challenge. By reducing drug particle size, nanocrystals increase dissolution rate and apparent solubility while avoiding the low payloads typical of conventional carrier-based systems, since the particles are composed predominantly of the active pharmaceutical ingredient [24]. This approach is particularly attractive for hydrophobic compounds whose preclinical or clinical development is limited by poor aqueous solubility. Surface engineering can further improve the performance of nanocrystal formulations. Coating drug nanocrystals with human serum albumin may enhance colloidal stability and in vivo behavior, while exploiting the biocompatibility, prolonged circulation and tumor-association properties of albumin [24–26]. We therefore reasoned that a carrier-free, albumin-coated nanocrystal formulation could convert VS1 into a deliverable agent and enable a necessary in vivo test of STARD3 inhibition as a fluoropyrimidine-potentiating strategy.

Here, we combine formulation science with structural biology, molecular modelling and functional pharmacology to develop STARD3 inhibition into a deliverable combination approach for CRC. We engineer an albumin-coated nanocrystal formulation that overcomes the poor solubility of VS1 and characterizes its size, morphology, crystallinity, molecular integrity, apparent solubility and release behavior. We also define the binding mode of VS1 by crystallography and molecular dynamics and show that albumin-enabled delivery of VS1 preserves antiproliferative activity and potentiates 5-FU in vitro and in vivo. Together, this work provides a delivery-enabled framework for advancing an otherwise undeliverable STARD3 inhibitor and supports albumin-coated nanocrystals as a platform for translating poorly soluble chemosensitizing agents into preclinical cancer therapy.

## MATERIALS AND METHODS

### Animal ethics

All procedures were conducted in accordance with EU Directive 2010/63/EU and the PREPARE, ARRIVE and OBSERVE guidelines and were approved by the National Ethical Committee and Ministry of Agriculture, Forestry and Food of the Republic of Slovenia (permission number: U34401-14/2023/13). Mice from Envigo were housed in filtertop cages (2–5 per cage) with bedding and enrichment materials and under controlled conditions of temperature (22 ± 2 °C), humidity (55 ± 10%), and light (12 h/12 h light–dark cycle) with unlimited access to food and water.

### Expression and purification of StARD3 START domain

The Lutein Binding domain of StARD3 (StARD3_LBD_, Lutein Binding Domain, residues 216-444) [27] was cloned into a pETDuet plasmid to express a protein with an uncleavable N-terminal His tag. The protein was expressed in *E. coli* BL21 StarTM (DE3) strain (Novagen, Merck, MA, US) in LB media. Cells were grown at 37°C to an OD600 of 0.7 and then transferred to 22°C. Protein expression was induced by the addition of 1 mM IPTG. After 18h, cells were centrifuged at 3,500 g for 30 minutes at 4 °C and resuspended in Lysis Buffer (20 mM Tris-HCl pH 8, 150 mM NaCl, 5 mM imidazole) complemented with 0,1 mg/mL DNase I, 1 mM phenylmethanesulfonyl fluoride (PMSF) (Merck, MA, US), 10 µM Leupeptin (Merck, MA, US), 2 µg/mL Aprotinin (Merck, MA, US) and 1mM TCEP. After cell disruption with a Sonicator (Bandeline), the crude extract was centrifuged at 20,000 g for 1hr at 4°C and the supernatant was incubated with 1 mL of Ni-NTA resin (Qiagen, Germany) for 45 minutes at 4°C. The resin was then washed with 10 Column Volume (CV) of Lysis Buffer supplemented with 25 mM imidazole and the protein was eluted at 200 mM imidazole. Appropriate factions were pooled and the salt concentration was reduced to 50 mM NaCl. Further steps followed the purification protocol reported by Horvath et al., 2016 [28]. The solution was applied to an HiTrap SP cation exchange column (Cytiva, MA, US), pre-equilibrated with Cation Exchange Buffer (25 mM Hepes pH 6.5, 10 mM NaCl, 0.5 mM EDTA, 1 mM TCEP). The column was washed with 10 CV of Cation Exchange Buffer and the protein was eluted with a linear gradient over 40 CV to a concentration of 200 mM NaCl. Appropriate fractions were pooled, concentrated, and applied to an analytical Superdex 75 10/300GL size exclusion column, using the protein crystallization buffer (20 mM Tris pH 7.5, 150 mM NaCl, 2 mM DTT). Fractions enriched in StARD3_LBD_ protein were pooled and concentrated **(Supplementary Fig. S1)**.

### Protein crystallization, data collection and structure determination

Freshly purified StARD3_LBD_ was concentrated to 8 mg/ml and centrifuged to remove insoluble material before crystallization. To obtain the StARD3_LBD_-VS1 complex, protein was mixed with a twofold molar excess of VS1, and the mixture was incubated at 37°C for 1 hour before crystallization. Crystals of StARD3_LBD_-VS1 complex were obtained in 0.1 M CHES pH 9.5, 0.2 M lithium sulphate, 0.3-0.6 sodium/potassium tartrate, 0.01M DTT using the hanging-drop vapor diffusion method at 22°C. Rod-shaped crystals were harvested, mounted on cryo-loops and flash-cooled in liquid nitrogen in mother liquor with no cryo-protectant. Diffraction data were collected on the XRD2 beamline (11.2R) at Elettra - Sincrotrone Trieste (Italy) and processed using XDS [29]. Phases were determined by molecular replacement method using AutoMR within the Phenix software suite [30] with the crystal structure of the apo protein (PDB-ID 5I9J) as a template, followed by AutoBuild [31]. The resulting model was manually improved and rebuilt through iterative model building using COOT [32]. Structural refinement was subsequently carried out using autoBUSTER [33]. Figures were prepared using PyMOL (The PyMOL Molecular Graphics System, Version 3.0 Schrödinger, LLC).

### MD Simulations

Molecular dynamics (MD) simulations were performed using AMBER, version 24 [34], using the ff14SB force field. General Amber force field (GAFF2) parameters were used for the ligand, whose partial charges were assigned using the AM1-BCC method through the Antechamber suite of AMBER 24. The complex was placed at the center of a rectangular parallelepiped box and solvated with a 20 Å water cap, generated using TIP3P explicit solvent model. Chloride ions were added to neutralize the system, which was subjected to a two-stage energy minimization. In the first stage, a harmonic potential of 100 kcal/mol•Å^2^ was applied to all solute atoms, thus only minimizing the solvent molecules. In the second stage, 5000 cycles of steepest descent followed by conjugate gradient (CG) were used to minimize the whole system, until a convergence of 0.05 kcal/Å•mol, since the position restraint was applied only on the protein α carbons. The minimized complex was used as input structure for the MD simulations, which were run using Particle Mesh Ewald (PME) electrostatics, a cutoff of 10 Å for the non-bonded interactions and periodic boundary conditions. SHAKE algorithm was used to constrain all bonds involving hydrogen atoms. Initially, an MD step of 0.5 ns in which the temperature of the system was raised from 0 to 300 K was performed using constant-volume periodic boundary conditions. An equilibration step of constant pressure periodic boundary MD was run for 3 ns, keeping the temperature of the system at the constant value of 300 K with Langevin thermostat. A time step of 2.0 fs was used for these two MD steps. The system was further equilibrated through an additional 3-ns constant pressure MD run, incorporating the hydrogen mass repartition (HMR) scheme [35], which allowed to extend the time step to 4.0 fs. A final 500-ns constant pressure HMR MD simulation was thus performed with a 4.0-fs time step. Throughout all MD steps, the harmonic potential of 10 kcal/(mol·Å^2^) on the protein α-carbons was maintained.

### SMD Simulations

Steered molecular dynamics (SMD) simulations were performed using AMBER24 [34] following a previously applied protocol [36]. The final MD frame obtained from the simulation of reference **VS1**-STARD3 complex was used as the starting structure for the SMD simulations, which were thus performed at the same constant-pressure periodic boundary conditions and at the constant temperature of 300 K. However, no position restraint was applied on the protein α carbons, and no HMR scheme was used during these simulations, which were thus performed employing a time step of 2.0 fs. Six different SMD simulations using six different initial pulling directions were performed. Each pulling direction was imposed by selecting a couple of atoms, one belonging to the protein and one belonging to the ligand, which set the initial distance to be stretched during the SMD. In all simulations, the atom of the ligand to which the pulling force was applied was C19, corresponding to the terminal carbon of the ligand phenyl ring. The protein atom from which the ligand was pulled in the six SMD simulations, respectively, corresponded to 1) the α carbon of I353, 2) the α carbon of S362, 3) the α carbon of G364, 4) the α carbon of G386, 5) the α carbon of F388, 6) the α carbon of W404. In all simulations, the ligand was pulled out from the protein binding site by increasing the initial distance between the selected couple of atoms by 30 Å, through the application of a spring constant of 5 kcal/mol•Å^2^. Each SMD simulation was performed at the constant velocity of 0.1 Å/ns by setting the simulation length to 300 ns; this allowed us to consider the simulated ligand unbinding process as reversible, and the pulling work associated to the process as the exact free energy [37,38]. Trajectory analyses including root-mean-square deviation (RMSD), root-mean-square fluctuation (RMSF) and generation of average ligand-protein structures were performed using the Cpptraj program [39] implemented in AMBER 24.

### Binding Energy Evaluations

All ligand-protein binding free energy evaluations were performed with AMBER 24 using the MM-GBSA method [40]. The trajectories corresponding to the last 200 ns of the classic MD simulation and the full trajectory of the selected SMD simulation (300 ns) were used for the evaluation, which was thus performed on a total of 200 and 300 MD frames (1 per ns), respectively. MOLSURF program and the MM-PBSA module of AMBER 24 were used to calculate nonpolar and polar energies, respectively, while van der Waals, electrostatic and internal contributions were estimated with SANDER module [41,42]. The ligand’s entropy was not considered in the calculation.

### VS1 nanocrystals production and albumin coating

The following raw materials were procured from Merck (Darmstadt, Germany): Pluronic® F-127 powder (F127), suitable for cell culture (Catalog No. P2443); hexadecyltrimethylammonium bromide (CTAB, ≥ 98%, Cat. No. H5882); human serum albumin (HSA) ≥ 96% (Catalog No. A3782); Dulbecco’s Phosphate Buffered Saline (DPBS) (Catalog No. D1408); deuterated dimethyl sulfoxide (DMSO-d6) (Catalog No. 175943); Methanol for LC/MS was sourced from Carlo Erba, Milan, Italy (Catalog No. A177150010), while chloroform (Catalog No. 67-66-3) was acquired from Thermo Fisher Scientific (Waltham, MA, USA). Water was purified using the Milli-Q® (Millipak® 0.22 μm) purification system. VS1 was produced by Enamine, Kiev, Ukraine. To create an aqueous dispersion of nanocrystals, the drug powder was processed using a method described in the literature by Park et al. [24] The described protocol initially involves producing a thin amorphous film by evaporating 24 mg of F127 and 6 mg of the drug (drug:F127 1:4) dissolved in 3 ml of chloroform using a rotary evaporator. The resulting thin film was then hydrated by introducing 6 ml of MQ water and subjected to sonication first with a bath sonicator (1 minute), followed by probe sonication (10 minutes, 40% amplitude) in an ice bath. This procedure produced a nanocrystal dispersion of VS1 stabilized with F127, designated as VS1-F. When 2.4 mg of CTAB was used in place of F127, the resulting dispersion - stabilized by CTAB - was referred to as VS1-C. To exchange surfactants with HSA, the drug-F complex was combined with HSA at a concentration of 4 mg/ml while rotating at 30 rpm, maintained at room temperature for 24 hours. The resultant mixture underwent centrifugation (38000 rpm, 4°C, 1 hour) twice to eliminate unbound ligands. The pellet derived from the purification process was then lyophilized and subsequently redispersed through probe sonication (10 minutes, 40% amplitude) in the original volume of MQ water. This ligand exchange procedure resulted in VS1-AF and VS1-AC.

### Spectrophotometric quantifications

Drugs were measured using UV-VIS spectrophotometry (Cary 100, Agilent) through a calibration curve at 270 nm. VS1-F was diluted in methanol for immediate analysis, while VS1-AF underwent centrifugation (2 minutes at 10000 rpm) prior to assessment. HSA levels were quantified using the Bradford assay, with absorbance readings taken at 595 nm based on a calibration curve. The concentration of the drug within nanocrystal dispersions was computed to determine the yield percentage employing the formula: Encapsulation efficiency (EE%) = [(experimentally determined drug concentration)/(weighted drug powder/MQ water volume)]*100. EE% reflects the efficiency of drug encapsulation within nanocrystals.

### Chemi-physical characterization

The properties of the VS1 molecule were predicted *in silico* using MolBook Pro software [43,44]. Additionally, the water solubility was experimentally assessed by dissolving 1 mg/ml of the inhibitor in MQ water, followed by mixing for 1 hour and centrifugation at 10000 rpm for 20 minutes. The resulting supernatant was subsequently analyzed to quantify the inhibitor, as previously documented for VS1-F.

### TEM, DLS, Z potential and XRD analyses

A solution droplet (approximately 25 μL) was applied to a 400-mesh holey film grid; following a 2-minute staining with 1% uranyl acetate, the sample was examined using a FEI Tecnai G2 transmission electron microscope operating at 100 kV (Hillsboro, Oregon, USA). Images were captured using a Veleta digital camera (Olympus Soft Imaging System). Image processing to delineate the crystals was performed with ImageJ (version 1.52a) software. Hydrodynamic diameter, polydispersity and zeta potential of the nanocrystals were assessed with a Zetasizer Nano particle analyzer (Malvern Panalytical, Malvern, UK). The nanocrystal solutions underwent an additional centrifugation step (1 hour, 4°C, 38000 rpm) to remove impurities, and the resulting pellets were subsequently freeze-dried. The acquired powders, along with other samples of interest, were evaluated using a Philips X-ray diffractometer equipped with a PW1050/70 goniometer. CuKα radiation (λ = 1.54178 Å) was employed, and diffractograms were recorded with a detection step of 0.05°. The intensity of the detected signals was measured as pulses per second. To facilitate comparisons among the spectra, they were normalized; it is important to note that the signal-to-noise ratio is affected by the amounts of samples used for the analysis.

### FT-IR and drug release tests

As previously mentioned for XRD analysis, the nanocrystals underwent additional purification, and their powdered form was obtained prior to conducting the FT-IR measurements. The relevant powders were utilized to fabricate KBr disks, and their FTIR spectra were recorded using a FTIR spectrophotometer (Perkin Elmer-Spectrum One).

A volume of 1 mL of inhibitor-F was introduced into a dialysis membrane Slide-A-Lyzer MINI Dialysis Device with a 20 k MWCO (Thermo Scientific, MA, USA) and subjected to dialysis against DPBS maintained at 37 °C while being mixed. Samples of the solution were taken at various time intervals, after which the inhibitor concentration was analyzed using UV-VIS spectrometry. The results are represented as a percentage of release, calculated as follows: Release % = 100 * ([Inh]t0 - [Inh]tx) / ([Inh]t0), where [Inh]t0 denotes the initial drug concentration in the inhibitor-F sample utilized for the experiment, and [Inh]tx refers to the inhibitor concentration measured in aliquots collected at different time intervals.

### Cell lines and Culture conditions

Colon cancer cell lines (HCT-116, COLO-201) and normal MRC-5 cells were purchased from ATCC, while HT-29 was purchased from CLS. HCT-116 and HT-29 were cultured in McCoy media (Gibco) containing 10% fetal bovine serum (FBS) (Gibco) and 1% penicillin/streptomycin (Gibco). COLO-201 cells were cultured in RPMI media (Gibco) containing 10% fetal bovine serum (FBS) (Gibco) and 1% penicillin/streptomycin (Gibco). Healthy MRC-5 cells were cultured in MEM media (Gibco) containing 1% non-essential amino acids (Gibco), 1% sodium pyruvate (Sigma), 10% FBS and 1% penicillin/streptomycin. Cells were incubated at 37 °C with 5% CO2 and 95% humidity. Mycoplasm contamination was checked monthly using MycoAlert® PLUS Mycoplasma Detection Kit (Lonza, LT07-705).

### Cell viability assay and synergism evaluation

Cell viability was assessed using the CellTiter-Glo® 2.0 Cell Viability Assay (Promega, WI, USA). Cells were treated with VS1, 5-fluorouracil, irinotecan, oxaliplatin and regorafenib (MedChemExpress, NJ, USA) at starting concentrations of 200, 100, 5, 5 and 10 µM, respectively, followed by 1:2 serial dilutions over 7 dose points; MRC-5 cells were treated with each drug starting from 200 µM. Luminescence readings were acquired after 96 h of treatment using a Tecan Infinite 1000 PRO plate reader (Tecan, Männedorf, Switzerland), following the manufacturer’s protocol. IC50 values were calculated by nonlinear regression with a dose-response inhibition model in GraphPad Prism 8.0.1 (GraphPad Software, La Jolla, CA, USA). Drug–drug interactions between VS1 and each chemotherapeutic were quantified by the Chou–Talalay combination index (CI) method [45], where cell viability data were converted to fraction affected (fa = 1 − viable fraction); the median-effect dose Dm and slope m for each single agent were derived from linear regression of log[fa/(1−fa)] versus log(D). The CI at each effect level was computed as CI = (D)₁/(Dx)₁ + (D)₂/(Dx)₂, where (D)ᵢ is the dose of drug i in combination and (Dx)ᵢ is the equieffective dose of drug i alone. CI < 1, CI = 1 and CI > 1 denote synergism, additivity and antagonism, respectively. All calculations and Fa–CI plots were generated with CompuSyn v.1.0 (ComboSyn Inc., Paramus, NJ, USA), after averaging Fa across n=3 biological replicates.

#### Reactive oxygen species measurements

Intracellular H₂O₂ was quantified with the ROS-Glo™ H₂O₂ Assay (Promega, WI, USA) in nonlytic mode, enabling ROS and viability readout from the same well. HCT-116 and COLO-201 cells seeded in 96-well plates (10000 cells/well) were treated with VS1, 5-FU or their combination, using sub-IC₅₀ concentrations applied at a fixed ratio in each cell line. The H₂O₂ Substrate (25 µM) was added at 18 h; at 24 h, 50 µL of supernatant was combined with 50 µL of ROS-Glo Detection Solution, incubated 20 min, and read on a Tecan Infinite 1000 PRO (Tecan, Männedorf, Switzerland). Viability was measured in parallel in the same wells with CellTiter-Glo® 2.0 (Promega). A positive control was also added, treating cells with exogenous H₂O₂ at 200 µM for one hour. The ROS signal was normalized to viability and expressed as fold change versus non-treated cells. Experiments were performed in biological triplicates.

#### In vivo experiments and histological analyses

HCT-116 cell line was cultured, harvested after trypsinization, and suspended in a saline solution enriched with 30% Matrigel HC (Corning^®^) to a density of 5*10^7^ cells/mL. Subsequently, 100 µL of this suspension was subcutaneously injected into the flank of mice. Upon reaching a volume of around 50 mm^3^, tumor-bearing mice were assorted into different groups: five treatment conditions were evaluated: control (CTR), VS1-F, VS1-AF, 5-FU and VS1-AF+5-FU.

Tumors were measured three times weekly using a digital Vernier caliper in three perpendicular directions (a, b, c), and volumes calculated using the formula V = a × b × c × π/6, where a, b, and c denote the diameters of the ellipsoid along its principal axes. To evaluate tumor growth delay, we identified the time it took for each tumor to double in size from its volume at the start of treatment. We then determined the tumor growth delay for each tumor by deducting its individual tumor doubling time from the average doubling time of the control group’s tumors. The average tumor growth delay was calculated for each treatment group. Animal well-being was monitored during the experiment by weighing the animal and visual inspection. Each condition involved eight tumors in nude mice, which received three weekly injections over a 21-day period at a dosage of 20 mg/kg per condition. Mice bodyweight as a sign of systemic toxicity of the treatments were evaluated throughout the entire treatment period.

For histological examination, a number of organs (brain, heart, lungs, small intestine, liver, kidneys, ovaries, uterus, spleen, bone marrow, lymph node and subcutaneous mass) of 24 animals were fixed overnight in 4% formaldehyde, and paraffin embedded. Four µm sections were examined after staining with Hematoxylin and Eosin (H&E).

### Statistical analyses

Statistical analyses were performed using GraphPad Prism 8.0.1 (GraphPad Software, San Diego, CA, USA). Differences between more than two groups were assessed by one-way analysis of variance (ANOVA) followed by Tukey’s post-hoc test for multiple comparisons, or by two-way ANOVA followed by Tukey’s post-hoc test where two independent variables were analyzed simultaneously. Data are expressed as mean ± SEM unless otherwise stated. A p-value < 0.05 was considered statistically significant and is indicated as *p < 0.05, **p < 0.01, ***p < 0.001, ****p < 0.0001.

## RESULTS

### Structure of VS1-bound form of StARD3_LBD_ protein

To define the molecular basis of STARD3 inhibition, we determined the crystal structure of the STARD3 ligand-binding domain (StARD3_LBD_) in complex with VS1. Crystals were obtained in the presence of a twofold molar excess of ligand, and the structure was solved at 2.1 Å resolution using the apo form as search model ([28], PDB ID: 5I9J). The refined model includes residues 230–444 and clear non-peptide density within the binding cavity corresponding to a bound VS1 molecule (**Supplementary Table S1**).

As previously described ([28,46]), StARD3_LBD_ adopts a helix-grip fold comprising a curved nine-stranded β-sheet and three α-helices, with the entrance to the cavity controlled by the Ω1 loop (**Fig. 1A**). Superposition with the apo structure showed that the overall fold is preserved upon ligand binding, while a local conformational rearrangement is observed at the Ω1 loop, consistent with ligand accommodation.

**Fig. 1.**
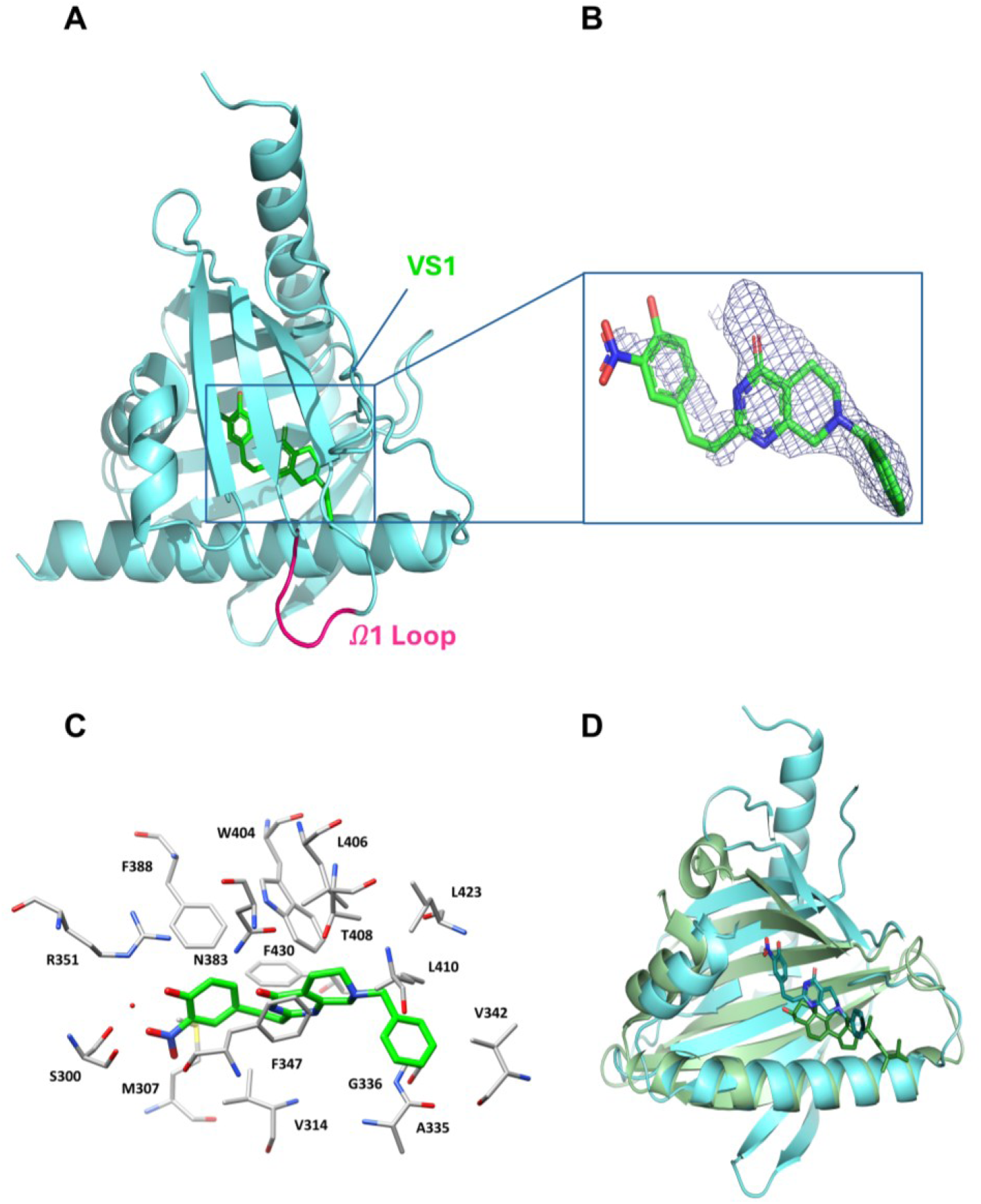
Crystal structure of the VS1-bound StARD3LBD domain. (**A**) Overall structure of the VS1-bound StARD3LBD domain in cartoon representation, with the Ω1 loop highlighted and VS1 shown as sticks. (**B**) Electron density map corresponding to the bound inhibitor. (**C**) Binding mode of VS1 into the StARD3LBD domain. The ligand is shown in green, while the surrounding protein residues are shown in gray. The structural water molecule is represented as a red sphere. (**D**) Structural comparison with the ergosterol-bound Ysp2p homolog, showing that VS1 occupies a deeper region of the cavity than the physiological sterol ligand.

VS1 binds within the characteristic L-shaped cavity of StARD3_LBD_ and is stabilized by a combination of hydrogen-bonding and hydrophobic interactions. The ligand core is well defined in the electron density map, whereas the nitrophenol moiety appears more flexible and was conservatively modeled in a single conformation with 50% occupancy (**Fig. 1B; Supplementary Fig. S2**). At the bottom of the cavity, the o-nitrophenolic group interacts with residues S300, M307, V314, R351, F388 and F430, with the phenolic -OH directly hydrogen-bonded to R351 and additional stabilization likely mediated by a structural water molecule. The aromatic ring also forms a T-shaped interaction with F338 and hydrophobic contacts with M307 and V314. The bicyclic core occupies the central portion of the cavity and interacts with W404, F347, L406 and T408, whereas the terminal benzyl group is positioned between the Ω1 loop and α4 helix, contacting A335, G336, V342, L423 and L410 (**Fig. 1C**). To assess whether VS1 competes with sterol binding, we compared the complex with the structure of the homologous StART-like domain of S. cerevisiae Ysp2p in complex with ergosterol (PDB ID: 6CAY). Unlike ergosterol, which binds near the cavity entrance, VS1 occupies a deeper region of the human STARD3 pocket, supporting a binding mode compatible with inhibition of cholesterol recognition and trafficking (**Fig.1D**).

### Molecular dynamics define a stable binding mode and a gated unbinding pathway for VS1

To further characterize the inhibitory mechanism of VS1, we performed molecular dynamics (MD) simulations on the VS1–STARD3 complex. Over >500 ns of classical MD in explicit solvent, the ligand remained stably bound within the STARD3 cavity, with an average RMSD of 0.9 Å relative to the starting pose, indicating minimal deviation from the crystallographic binding mode (**Supplementary Fig. S3**). All major ligand moieties preserved their orientation within the binding pocket, supporting a structurally stable bound state.

We next used steered molecular dynamics (SMD) simulations to investigate the dissociation pathway of VS1 and to identify transient conformational states that may inform future structure–activity relationship studies. Because access to the ligand-binding cavity is controlled by the Ω1 loop, we focused on ligand egress through this region. Six distinct unbinding trajectories (P1–P6) were simulated using different pulling directions defined by the Cα atoms of residues I353, S362, G364, G386, F388 and W404, respectively (**Supplementary Fig. S4A**). In each case, the distance between the selected residue and the terminal atom of the ligand phenyl ring was increased by 30 Å over 300 ns at a constant pulling velocity of 0.1 Å/ns (see Methods). Under these conditions, the simulated unbinding process could be treated as quasi-reversible, allowing the associated work to approximate the free-energy variation of the process [37,38].

Among the six trajectories, pathway P2 was the most energetically favorable, with a total free-energy cost of 22.5 kcal/mol, whereas the other pathways were associated with substantially higher values, ranging from 30.3 kcal/mol to 40.0 kcal/mol (**Supplementary Fig. S4B**). We therefore selected P2 for detailed structural and energetic analysis.

Along this pathway, VS1 progressively exited the binding cavity during the first ∼170 ns of SMD (**Fig. 2A**). The ligand began to protrude from the pocket after ∼50 ns, was largely displaced by ∼150 ns, and was fully outside the cavity by ∼170 ns, after which it detached from the protein surface and diffused into bulk solvent. This progression was reflected in the ligand RMSD profile, which increased gradually during the early phase of dissociation and rose sharply after complete exit from the binding site (**Fig. 2A**).

**Fig. 2.**
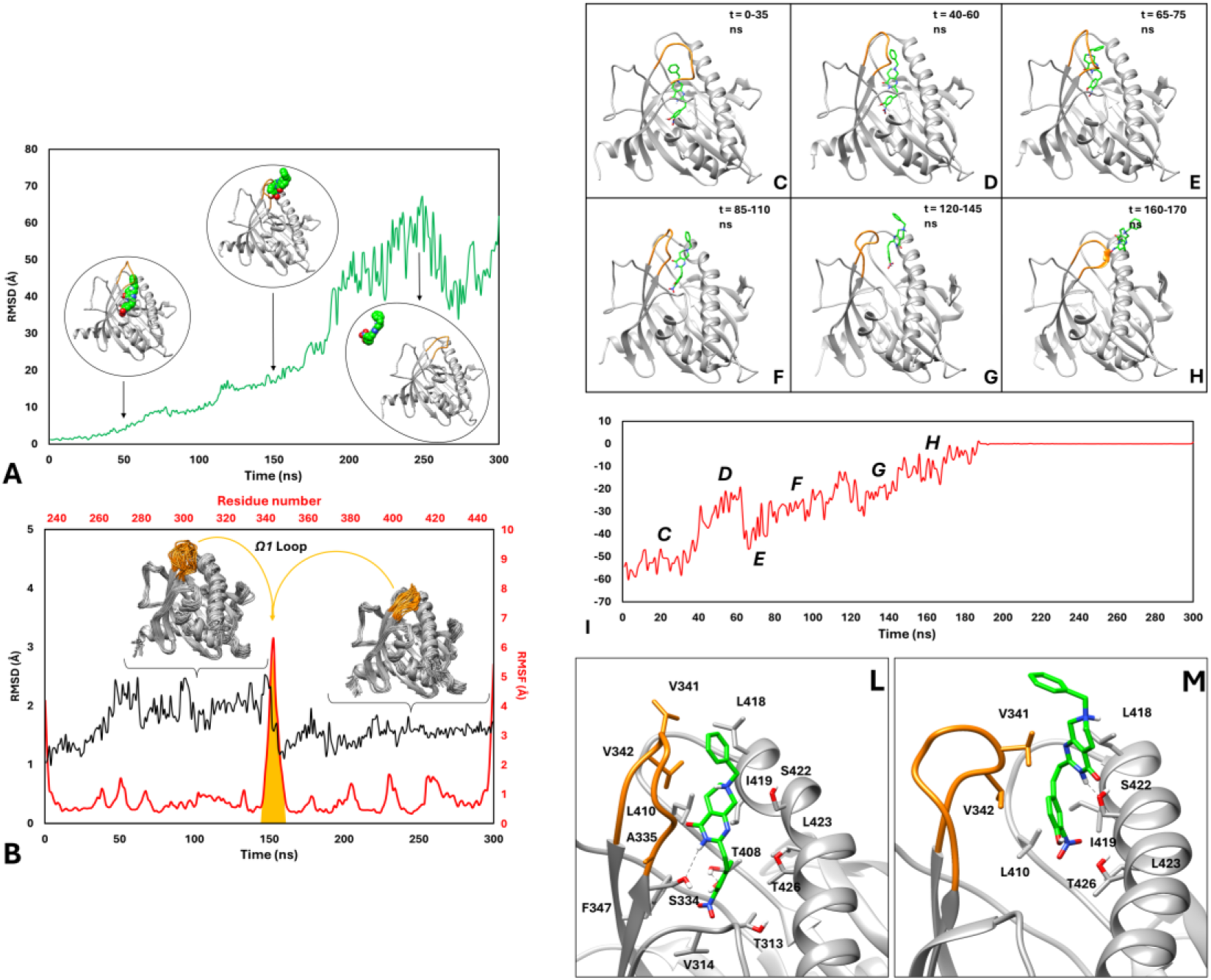
Ligand dissociation from STARD3 is mediated by transient opening of the Ω1 loop. (**A**) RMSD of VS1 during the selected SMD trajectory relative to the starting coordinates. Representative conformations at 50, 150 and 250 ns are shown. VS1 is shown as green spheres; STARD3 is shown as grey ribbons, with the Ω1 loop highlighted in orange. (**B**) Protein conformational dynamics during ligand dissociation. Black line, Cα RMSD of STARD3 during SMD; red line, residue-wise Cα RMSF. Representative protein conformational ensembles from 100–150 ns and 180–300 ns are shown. The Ω1 loop is highlighted in orange. **Energetic landscape and metastable intermediates of VS1 unbinding from STARD3**. (**C-H)** Average structures of the VS1–STARD3 complex at distinct stages of the selected SMD trajectory: a, 0–35 ns; b, 40–60 ns; c, 65–75 ns; d, 85–110 ns; e, 120–145 ns; f, 160–170 ns. VS1 is shown as green sticks and STARD3 as grey ribbons, with the Ω1 loop highlighted in orange. (**I**) Per-frame MM-GBSA binding free-energy profile along the SMD trajectory. Labels C–H indicate the intervals corresponding to the average conformations shown in panels C–H. (**L-M**) Detailed views of the two metastable intermediate binding modes observed at **L**, 85–110 ns and **M**, 120–145 ns. Hydrogen bonds are shown as black dashed lines.

Despite ligand dissociation, the overall conformation of STARD3 remained stable throughout the simulation. The average Cα RMSD was 1.7 Å, indicating preservation of the global fold (**Fig. 2B**). By contrast, residue-level fluctuation analysis revealed pronounced mobility in the Ω1 loop, with RMSF values peaking at residues 335–345 and reaching up to 6.3 Å, whereas the remainder of the protein fluctuated around ∼1 Å on average (**Fig. 2B**). These data indicate that ligand egress is mediated by a localized opening of the Ω1 loop, which transiently adopts an open conformation to permit ligand passage and subsequently returns toward its native state once dissociation is complete.

To quantify the energetic evolution of the unbinding process, we performed MM-GBSA calculations on snapshots extracted from both the classical MD trajectory and the selected SMD trajectory. The average binding free energy of the equilibrated bound complex, calculated from the last 200 ns of classical MD, was −52.0 kcal/mol, consistent with a stable ligand–protein interaction. Per-frame MM-GBSA analysis along the SMD trajectory revealed a progressive loss of binding energy during dissociation, accompanied by the appearance of transient metastable states (**Fig. 2C–I**). During the first 35 ns of SMD, VS1 retained a binding mode closely resembling the crystallographic pose, with the Ω1 loop in a closed conformation and an average binding free energy of −52.8 kcal/mol (**Fig. 2C,I**). Between 40 and 60 ns, the benzyl moiety began to push against the Ω1 loop and protrude toward the cavity entrance, resulting in a marked reduction in binding energy to −29.3 kcal/mol (**Fig. 2D,I**). A transient recovery in interaction energy was observed between 65 and 75 ns, when the ligand established new contacts with residues of the opened Ω1 loop, yielding an average binding free energy of approximately −39 kcal/mol (**Fig. 2E,I**). Thereafter, binding energy progressively decreased as the ligand became increasingly solvent-exposed and lost stabilizing contacts within the pocket.

Two metastable intermediate states were particularly prominent during the dissociation trajectory. In the first, observed between 85 and 110 ns, VS1 remained partially anchored at the cavity entrance despite substantial solvent exposure (**Fig. 2F,L**). In this conformation, the phenolic OH formed a hydrogen bond with T408, while the bicyclic core established an additional hydrogen bond with S334 and hydrophobic contacts with V341, V342 and I419. In the second intermediate, observed between 120 and 145 ns, the benzyl fragment was fully solvent-exposed, whereas the bicyclic core remained stabilized between the Ω1 loop and the α4 helix through hydrophobic interactions with V341 and L418 and a hydrogen bond with S442 (**Fig. 2G,M**). The binding free energies of these two states differed by less than 4 kcal/mol (**Fig. 2I**), indicating that both represent energetically relevant intermediates along the preferred exit pathway.

Notably, comparison of these two conformations suggests that the bicyclic core of VS1 undergoes a ∼180° reorientation during dissociation, accompanied by a switch in anchoring interactions from S334 to S442. This transition likely facilitates ligand passage through the cavity entrance and points to a functional role for these residues in guiding both ligand binding and release. Together, these simulations identify the Ω1 loop as a dynamic gate controlling ligand egress and reveal metastable binding states that may inform the design of next-generation STARD3 inhibitors.

### Physicochemical properties of VS1 and nanocrystal formulation

VS1, chemically defined as 2-[(E)-2-(4-hydroxy-3-nitrophenyl)ethenyl]-7-benzyl-5,6,7,8-tetrahydro-1,3,7-triaza-4(1H)-naphthalenone, exhibits moderate lipophilicity, with a predicted LogP value of 3.52. In silico estimations suggested limited aqueous solubility, with a predicted value of approximately 63 µg mL⁻¹. Experimental measurements confirmed the poor water solubility of VS1, revealing a solubility below 10 µg mL⁻¹ in milli-q water. This low aqueous solubility represents a major limitation for biological studies and future translational applications. To overcome this constraint, VS1 was formulated as nanocrystals stabilized either with Pluronic F127 (VS1-F) or with human serum albumin following surfactant exchange (VS1-AF). Both formulations generated stable aqueous dispersions of elongated nanocrystals, enabling quantitative assessment of particle size and morphology. Dynamic light scattering showed hydrodynamic diameters in the nanometric range and polydispersity indices of approximately 0.3 for all formulations, indicating acceptable colloidal homogeneity for anisotropic particles (**Fig. 3A**). Quantification of drug loading confirmed high encapsulation efficiency (EE%) for VS1-F, which was largely preserved following surfactant-to-albumin exchange in VS1-AF (**Fig. 3B**). Transmission electron microscopy further showed that nanocrystallization markedly reduced both particle length and diameter relative to the free compound, yielding smaller and more uniform rod-shaped particles in both VS1-F and VS1-AF preparations (**Fig. 3C,D; Supplementary Fig. S5**). Occasional aggregation in VS1-AF was attributed to drying artifacts, whereas the fragmentation observed in both nanocrystal formulations was likely due to sonication. The crystalline nature of the formulated particles was confirmed by X-ray diffraction, which showed that both VS1-F and VS1-AF retained a solid crystalline organization after nanocrystallization (**Supplementary Fig. S6**). Although the crystallization process altered the crystal phase relative to the free compound, the available data did not allow us to determine whether albumin exchange induced additional structural rearrangements because of the limited signal-to-noise ratio. FTIR analysis revealed that the spectra of VS1, VS1-F, and VS1-AF are nearly identical, showing only minor shifts in regions associated with water-related vibrational modes rather than differences stemming from the crystalline phase (**Fig. 3E; Supplementary Table S2**). These results indicate that the formulation process preserves the structural integrity of the inhibitor molecule. We next assessed drug release behavior under physiologically relevant conditions. VS1-F displayed a sustained release profile in DPBS at 37 °C, consistent with gradual drug liberation from a crystalline matrix (**Fig. 3F**). Notably, nanocrystallization increased the apparent aqueous solubility of VS1 by more than 14-fold relative to its predicted intrinsic solubility at neutral pH, supporting efficient drug incorporation and improved dispersibility. Additional CTAB-based nanocrystal formulations were generated as formulation controls. VS1-C showed high encapsulation efficiency, whereas albumin functionalization of this system (VS1-AC) resulted in a modest reduction in drug loading (**Supplementary Table S3**). FTIR spectra and peaks assignments of VS1-AC are displayed in **Supplementary Fig. S7** and **Supplementary Table S4**. Together, these results show that nanocrystal engineering substantially improves the physicochemical properties of VS1 and supports the use of albumin-coated formulations for downstream biological evaluation.

**Fig. 3.**
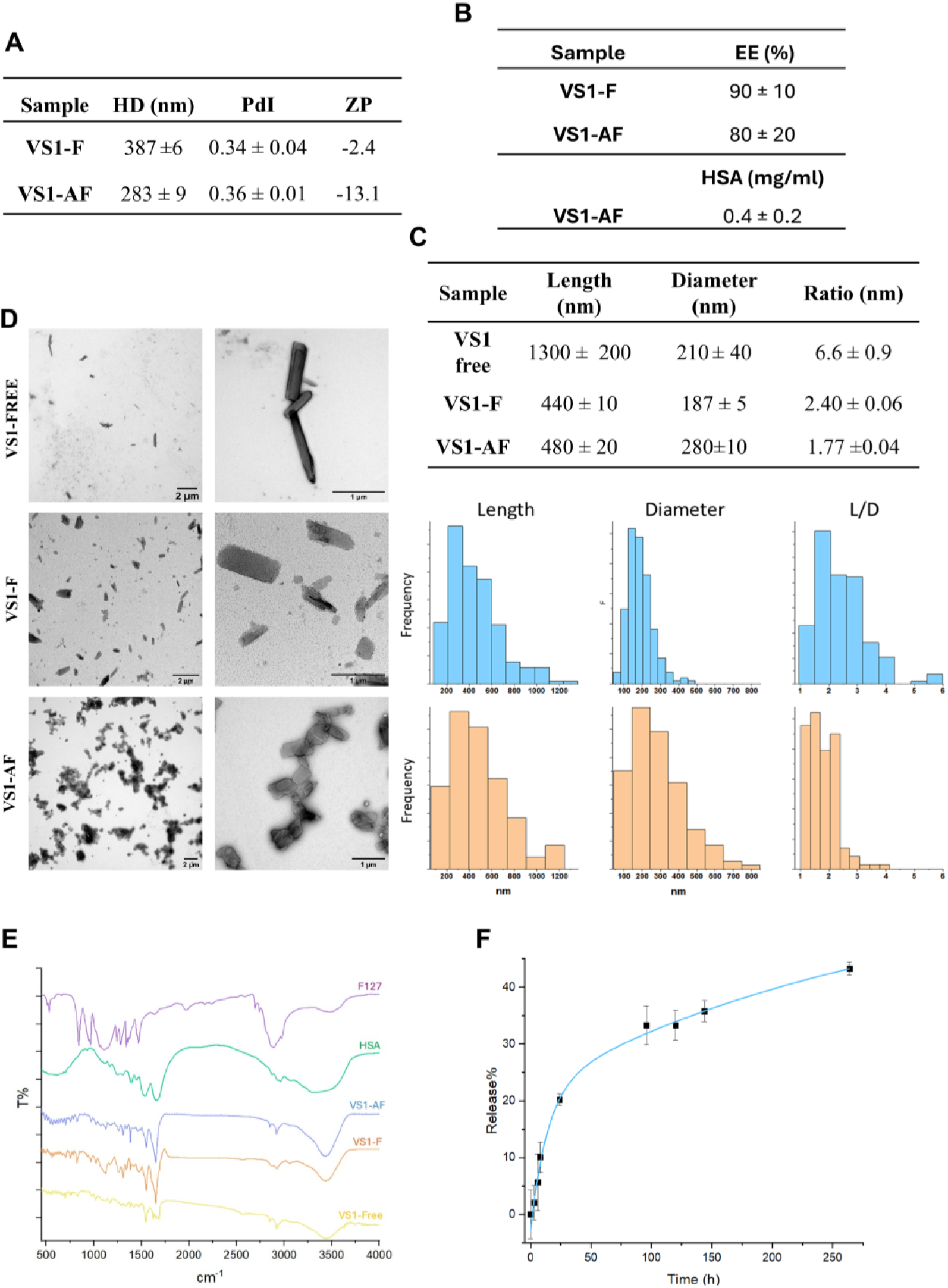
Nanocrystal formulation improves the physicochemical properties of VS1. **(A)** Dynamic light scattering analysis showing hydrodynamic diameter, polydispersity index and zeta potential of the indicated formulations. **(B)** EE% of VS1 nanocrystals and HSA quantified in VS1-AF. (**C**) Quantification of nanocrystal dimensions from TEM images. **(D)** Representative TEM images of VS1-free, VS1-F and VS1-AF preparations. **(E)** FTIR spectra of F127, HSA, free VS1, VS1-F and VS1-AF. **(F)** In vitro release profile of VS1 from VS1-F in DPBS at 37 °C.

### VS1 synergizes with 5-FU in HCT-116 and partially in HT-29 colon cancer cell lines

VS1 cytotoxicity was tested in HCT-116, HT-29 and COLO-201 colorectal cancer cell lines, alongside with normal MRC-5 fibroblasts. VS1 displayed IC50 values in the 23.6–58.5 µM range across the CRC panel, with the highest sensitivity observed in COLO-201 (**Table 1**).

**Table 1:** IC_50_ on colon cancer cell lines with VS1. The activity was evaluated in serial dilutions starting from different concentrations (VS1 - 200 µM; 5-FU - 100 µM; Irinotecan - 5 µM; Oxaliplatin - 5 µM; Regorafenib - 10 µM). IC_50_ was measured with CellTiter-Glo® 2.0 Cell Viability Assay. Experiments were conducted in biological triplicates.

| IC<sub>50</sub> (μM) | VS1 | Regorafenib | 5-Fluorouracil | Oxaliplatin | Irinotecan |
| --- | --- | --- | --- | --- | --- |
| **HCT-116** | 58±32 | 1.5±1 | 3.6±1.8 | 0.5±0.3 | 0.9±0.4 |
| **HT-29** | 55±11 | 2.4±0.2 | 2.3±1.5 | 0.4±0.03 | 3.1±1 |
| **COLO-201** | 23±7.8 | 3.8±1.5 | 8.8±4.5 | 1.1±0.5 | 2.1±0.7 |
| **MRC5** | >200 | 11.6±3 | >200 | >200 | 25±6.9 |

Importantly, the IC_50_ in MRC-5 cells exceeded 200 µM, indicating minimal cytotoxicity toward normal cells. To evaluate whether STARD3 inhibition could potentiate standard chemotherapeutic agents used in CRC treatment, VS1 was tested in combination with 5-fluorouracil (5-FU), oxaliplatin, irinotecan, or regorafenib in three CRC cell lines representing distinct molecular backgrounds: HCT-116 (MSI-H, KRAS-mutant, p53 wt), HT-29 (MSS, BRAF V600E, p53-mutated), and COLO-201 (MSS, BRAFV600E, derived from a 5-FU-pretreated patient). Combinations were assessed using a fixed-ratio diagonal design with serial 1:2 dilutions, performed in technical triplicates across n = 3 independent biological experiments (concentration ranges in **Supplementary Table S5**). Drug interactions were first quantified by the Chou-Talalay method, which yields Combination Index (CI) values at defined effect levels — ED_50_, ED_75_, and ED_90_ — corresponding to 50%, 75%, and 90% growth inhibition (CI < 1, synergism; CI ≈ 1, additivity; CI > 1, antagonism). VS1 + 5-FU showed strong cell-line specificity (**Fig. 4A**): HCT-116 displayed consistent synergism across all effect levels (CI = 0.82-0.86), and HT-29 was synergistic at intermediate effects (CI = 0.82 at ED_50_ and 0.77 at ED_75_), while COLO-201 was consistently antagonistic (CI = 1.61 at ED_50_, 1.62 at ED_75_). ED_90_ estimates for HT-29 and COLO-201 were not reported, as the 5-FU dose required exceeded the pharmacologically relevant range (>100 µM), reflecting the shallow dose-response curves of these lines. Isobologram analysis corroborated this pattern, with the combination point falling below the line of additivity in HCT-116 and HT-29, and above it in COLO-201 (**Fig. 4B**). The remaining VS1 + chemotherapy combinations did not produce consistent synergism in any line (VS1 + oxaliplatin: antagonistic across the panel; VS1 + regorafenib: largely additive; VS1 + irinotecan: additive to marginally synergistic in HCT-116, mixed in HT-29, antagonistic in COLO-201).

**Fig. 4:**
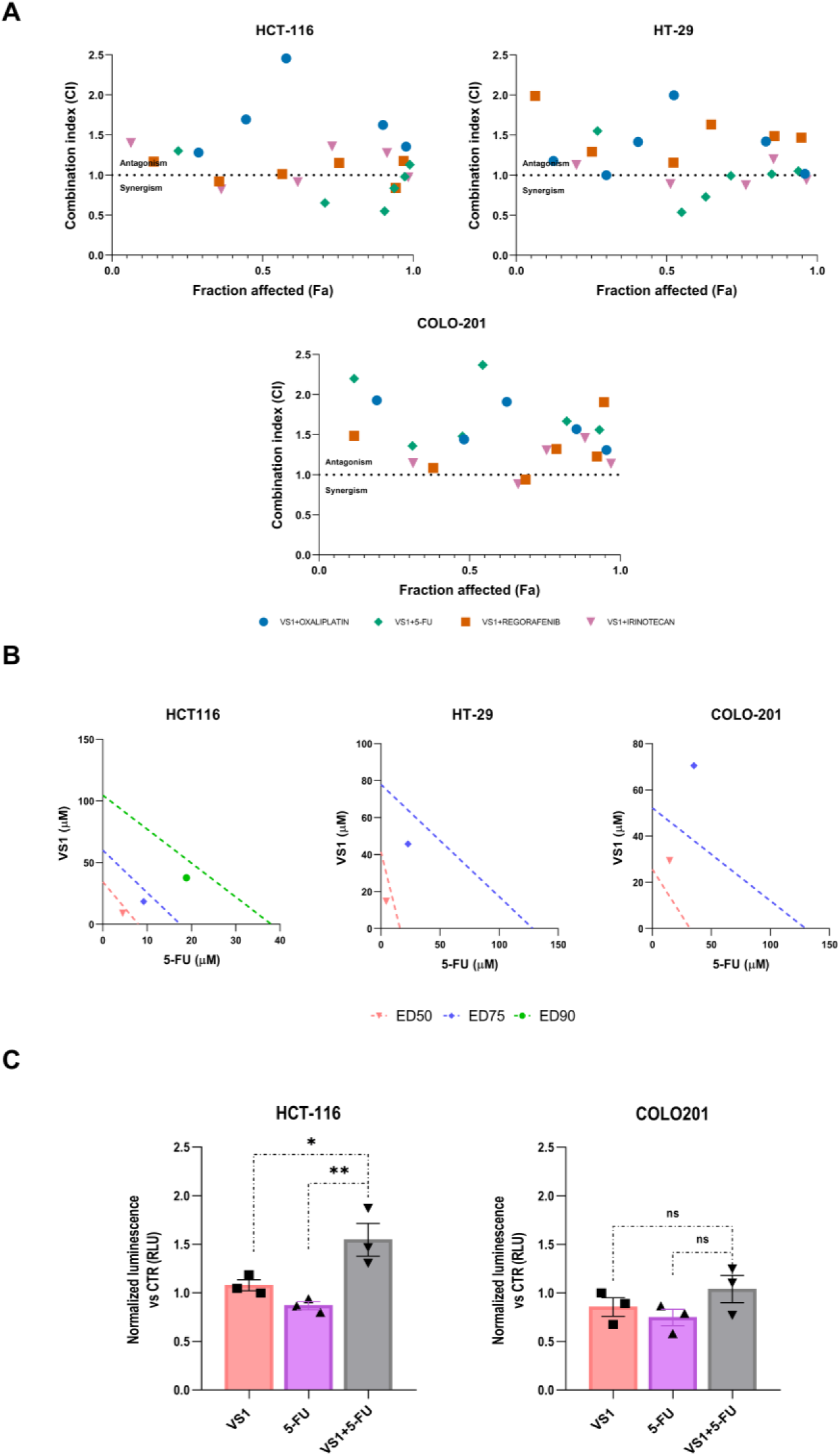
**(A) Combination Index (CI) plotted as a function of fraction affected (Fa) for each VS1-based combination, calculated according to the Chou-Talalay method.** Each symbol represents a single experimental dose combination. The dashed line at CI = 1 separates the synergistic (CI < 1) from the antagonistic region (CI > 1). Chou-Talalay analysis was performed with CompuSyn after averaging Fa across n=3 biological replicates. (**B**) **Isobologram analysis of VS1 + 5-FU at ED_50_, ED_75_, and ED_90_ (where applicable).** Dashed lines connect the equi-effective single-agent doses (line of additivity); experimental combination points falling below the line indicate synergism, while points above indicate antagonism. (**C**) **Effect of VS1 and 5-FU co-treatment on intracellular ROS production**. ROS levels were evaluated via normalized luminescence (RLU) vs non-treated cells in HCT-116 and COLO-201 colorectal cancer cell lines following treatment with VS1, 5-FU, or their combination (VS1+5-FU). One-way ANOVA followed by Tukey’s multiple comparison test, *p < 0.05; **p < 0.01; ***p < 0.001. Data are expressed as mean ± SEM.

Quantitative analysis of effective doses further supported these findings (**Table 2**): in HCT-116, the combination reduced the 5-FU dose required at ED_50_ by 1.8-fold (from 8.0 to 4.5 µM) and the VS1 dose by 3.8-fold (from 34.3 to 9.0 µM), with comparable dose-sparing in HT-29 (5-FU 2.2-fold, VS1 2.8-fold). In contrast, COLO-201 showed only modest reductions in the 5-FU dose within the combination (DRI ≈ 2.2-3.7), insufficient to overcome the antagonistic interaction, and no VS1 dose-sparing (DRI < 1). Although COLO-201 cells retain a relatively low IC_50_ for 5-FU, their shallow dose-response curves (Hill slope m = 0.78) point to pronounced response heterogeneity, likely reflecting traits established in vivo prior to isolation from ascites. Overall, Chou-Talalay analysis identified VS1+5-FU as the only combination showing reproducible synergy across the panel, robust and consistent across effect levels in MSI/KRAS-mutant HCT-116, partial and dose-restricted in MSS/BRAF-mutant HT-29, and antagonistic in MSS/BRAF-mutant COLO-201. This molecularly stratified pattern of activity, together with the established central role of 5-FU in first-line CRC regimens, identified VS1 + 5-FU in HCT-116 as the lead combination for in vivo validation.

**Table 2:**
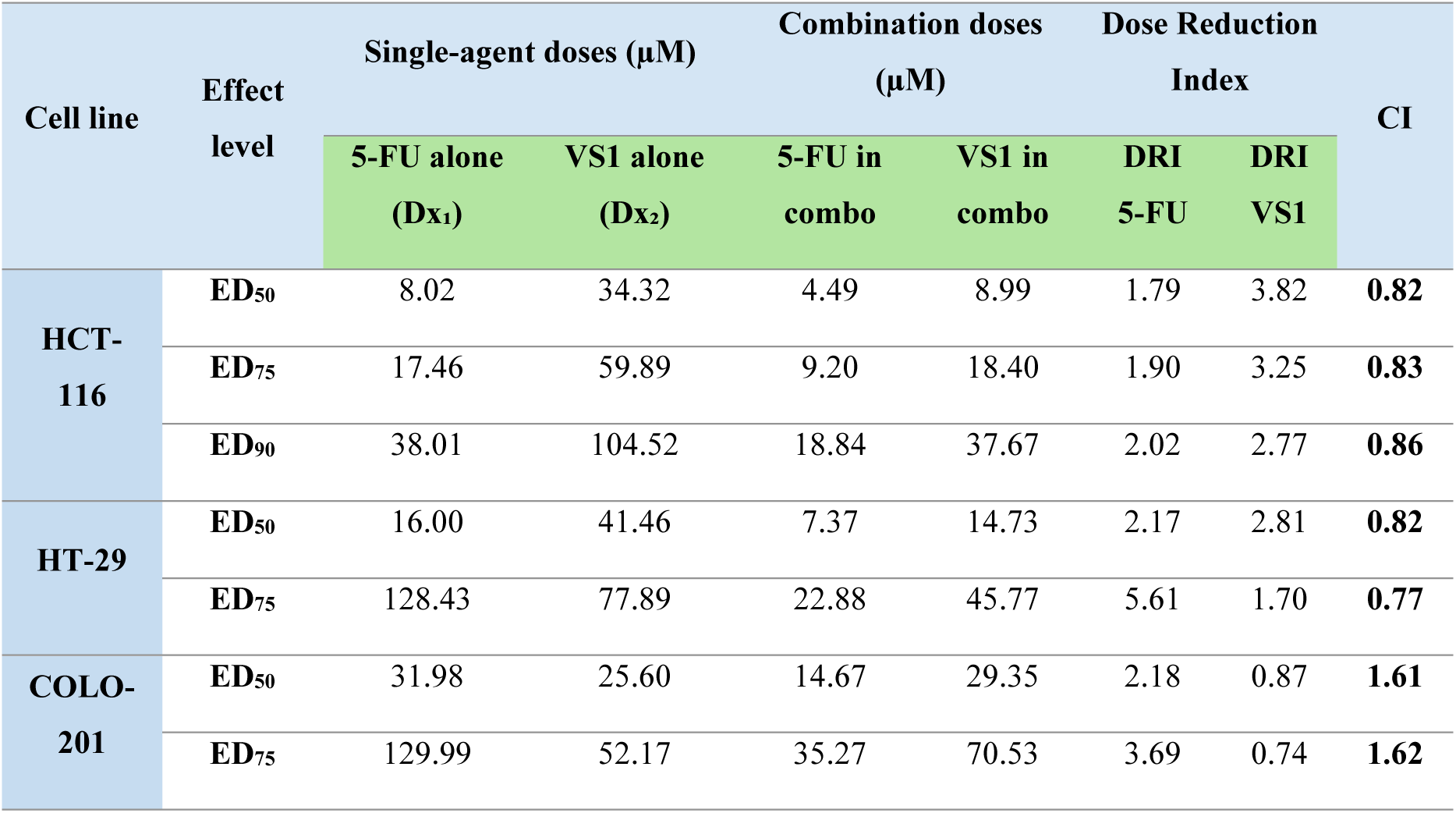
Combination Index values were calculated according to the Chou-Talalay method at ED_50_, ED_75_, and ED_90_ (CI < 1, synergism; CI ≈ 1, additivity; CI > 1, antagonism) . Dx values represent the single-agent doses estimated by the median-effect equation to produce the indicated effect level, while "5-FU in combo" and "VS1 in combo" represent the dose of each drug within the combination required to reach the same effect. The Dose Reduction Index (DRI) is defined as Dx/dose in combination, with DRI > 1 indicating dose sparing. Note that Dx values derived from the median-effect equation may differ from IC_50_ values obtained by three-parameter logistic regression (Table 1), as the two models exclude or include different data points at the extremes of the dose-response curve. ED_90_ doses for 5-FU exceeded the pharmacologically relevant range (>100 µM) in HT-29 and COLO-201, reflecting the shallow dose-response curves of these lines and the resulting extrapolation by the median-effect fit.

### The VS1 and 5-FU Combination Triggers Cell Line-Specific Synergistic ROS Accumulation

Treatment with the VS1 + 5-FU combination induced a significant increase in ROS production compared to either single agent in HCT-116, whereas no such cooperative effect was observed in COLO-201, where the combination did not differ significantly from VS1 or 5-FU alone (**Fig. 4C**; concentrations in **Supplementary Table S6**). This oxidative amplification specifically in HCT-116 supports a pro-oxidant component of the VS1 + 5-FU synergy, whereby VS1 sensitizes cells to 5-FU-mediated cytotoxicity by overwhelming their antioxidant defenses. The absence of the same effect in COLO-201 indicates that this cooperative ROS accumulation is context-dependent and shaped by the distinct genetic and metabolic background of each cell line, paralleling the cell-line-specific pattern of cytotoxic synergy identified by Chou-Talalay analysis.

### Albumin-formulated VS1 enhances 5-FU efficacy in vivo

Before evaluating the combination in vivo, the albumin-formulated nanocrystal preparation (VS1-AF) was tested in combination with 5-FU in HCT-116 cells. VS1-F retained antiproliferative activity comparable to the free compound and showed a strong synergism with 5-FU across all effect levels (CI = 0.49, 0.45, and 0.41 at ED_50_, ED_75_, and ED_90_, respectively; **Supplementary Fig. S8A**). Isobologram analysis confirmed the marked dose-sparing effect of the combination (**Supplementary Fig. S8B**), supporting the use of the nanocrystal formulation for *in vivo* studies. We next assessed the therapeutic efficacy of the albumin-formulated nanocrystal preparation (VS1-AF) in combination with 5-FU in an HCT-116 xenograft model, which are schematically represent in **Fig. 5A**. Compared with either monotherapy, the combination treatment produced a markedly stronger antitumor effect, resulting in sustained suppression of tumor growth over time (**Fig. 5B**). Tumor growth kinetics analysis revealed that VS1-AF+5-FU combination significantly prolonged both tumor doubling time (DT) and tumor growth delay (TGD) compared to CTRL (∗p < 0.05; **Fig. 5C–D**). Single-agent treatments (VS1-F, VS1-AF, and 5-FU) showed a trend toward increased DT and positive TGD values but failed to reach statistical significance. Notably, the combination group displayed greater response consistency across individual animals, suggesting that VS1-AF sensitizes HCT-116 tumors to 5-FU beyond simple additive cytotoxicity. Histological analysis further showed that combination treatment increased the proportion of necrotic tissue within the tumor compared with controls, consistent with enhanced induction of tumor cell death (**Fig. 5E; Supplementary Fig. S9**). VS1-AF was selected for combination studies based on its improved formulation profile, including better tolerability and bioavailability relative to the non-albumin formulation. Treatment was generally well tolerated, as indicated by stable body weight throughout the 21-day treatment period (**Fig. 5F**). Histopathological examination revealed reversible liver alterations characterized by cloudy swelling of hepatocytes across treatment groups, but the combination treatment did not exacerbate the effect. No relevant pathological alterations were detected in the other organs examined (**Fig. 5G**). Together, these results show that albumin-enabled delivery of VS1 enhances the therapeutic efficacy of 5-FU in vivo without increasing systemic toxicity, supporting STARD3 inhibition as a combinatorial strategy to improve fluoropyrimidine-based treatment in colorectal cancer.

**Fig. 5:**
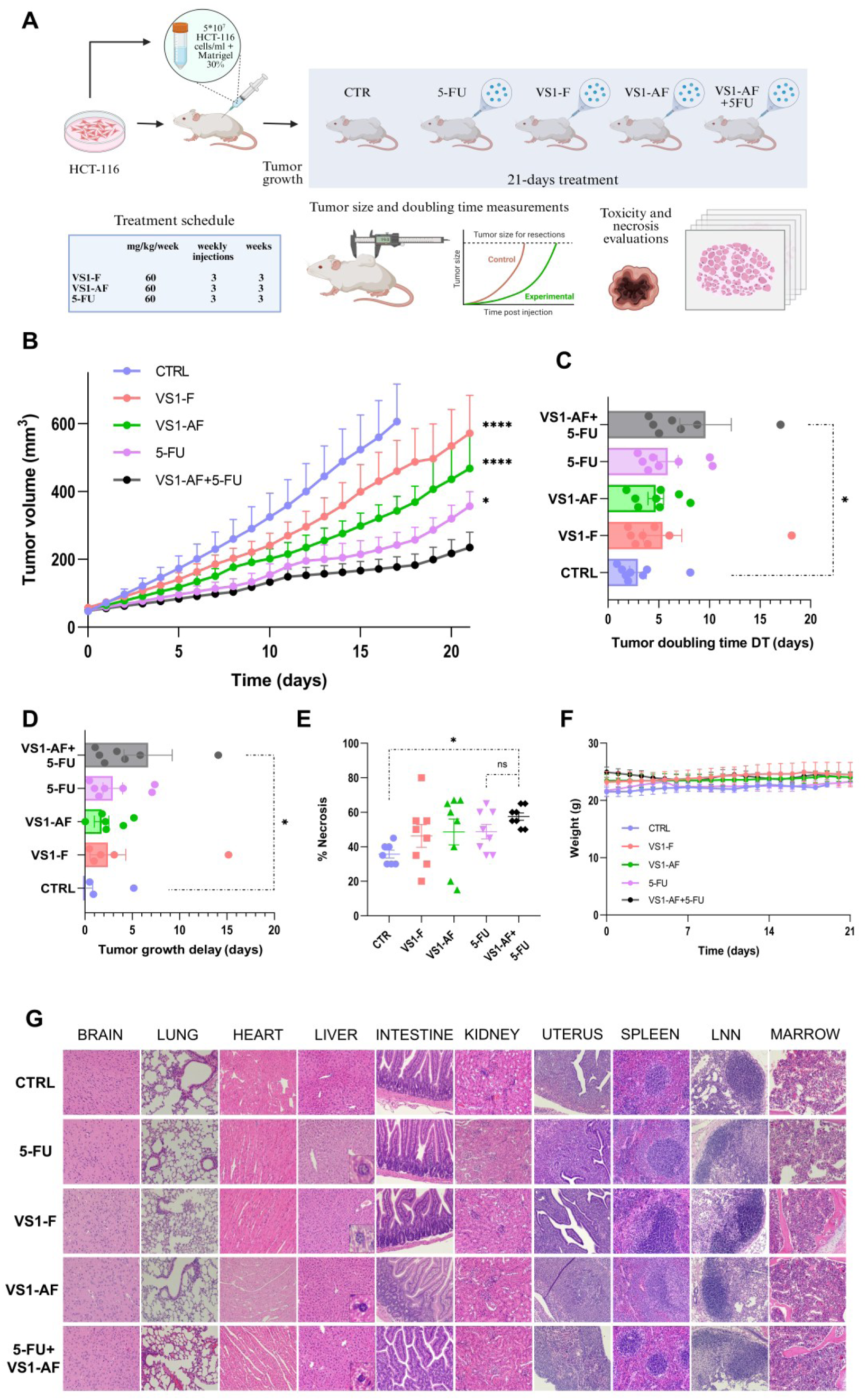
(**A**) **Schematic representation of in vivo experiments**. (**B**) **Tumor volume (mm^3^) evaluation after treatment.** The analysis highlights a greater efficacy in tumor reduction for drugs combinations. Asterisks indicate statistically significant differences compared to the VS1-AF+5-FU combination group. Two-way ANOVA followed by Tukey’s multiple comparison test. *p < 0.05; **p < 0.01; ***p < 0.001; ****p < 0.0001. (**C**) **Tumor doubling time (DT) analysis.** The combination treatment increase DT compared to the 5-FU-only group. One-way ANOVA followed by Tukey’s multiple comparison test, *p < 0.05. (**D**) **Tumor growth delay (TGD) across treatment groups**. VS1-AF+5-FU produced a statistically significant increase in TGD compared to CTRL. One-way ANOVA followed by Tukey’s multiple comparison test, *p < 0.05. (**E**) **Proportion (%) of tumor necrosis in cutaneous/subcutaneous tumor.** One-way ANOVA followed by Tukey’s multiple comparison test, *p < 0.05. (**F**) **Body weight analysis over the 21-period treatment for VS1 nanocrystals utilized in the test**. Drugs were well tolerated. (**G**) **Histological analysis**. Drug-induced liver alterations (cloudy swelling of hepatocytes) were detected across treatments (zoom in). Degenerated liver cells appear swelled, pale (cloudy) with poorly delineated and displaced nuclei.

## DISCUSSION

CRC remains a major cause of cancer-related mortality, and resistance to fluoropyrimidine-based regimens continues to limit therapeutic benefit in a substantial proportion of patients [1,5–7]. In this study, we identify STARD3 as a druggable vulnerability in CRC and provide the delivery platform needed to test it *in vivo*. We show that the small molecule VS1 binds directly to the STARD3 START domain, occupies the cholesterol-binding cavity, and, when reformulated as a carrier-free, albumin-coated nanocrystal, enhances the antitumor activity of 5-FU in vitro and in vivo. Because VS1 is essentially insoluble in water, this formulation step is not a technical adjunct but an enabling requirement for in vivo applicability.

Two advances emerge from this work. The first advance is the structural definition of STARD3 inhibition, which provides the rationale for the delivered payload. Although VS1 had previously been identified through in silico screening and shown to bind STARD3 with micromolar affinity [20,23], direct structural evidence was lacking. The crystallographic data reveal that the first in class inhibitor VS1 binds deeply within the STARD3 cavity, consistent with competitive interference with sterol recognition. Molecular dynamics further support a stable bound state and identify the Ω1 loop as a dynamic gate regulating ligand egress. The metastable states observed during dissociation suggest that residues within the Ω1 loop and the α4 region contribute to ligand stabilization and may be exploitable for future optimization, providing a structural framework for medicinal-chemistry efforts aimed at improving potency and residence time.

From a delivery standpoint, the second advance is the conversion of this structurally validated but undeliverable inhibitor into an in vivo-active agent using a simple, scalable nanocrystal approach. Drug nanocrystals are particularly suited to molecules such as VS1 because they enhance the dissolution rate and apparent solubility without the low drug payloads typical of carrier systems, as the particles are composed almost entirely of the active ingredient [24]. Here, nanocrystallization increased the apparent solubility of VS1 by more than an order of magnitude and produced a sustained, dissolution-controlled release profile under physiological conditions, while XRD and FTIR confirmed that crystallinity and molecular integrity were preserved. Surfactant exchange for human serum albumin yielded a carrier-free, albumin-coated nanocrystal that retained antiproliferative activity and improved tolerability relative to the non-coated formulation. Albumin is an attractive coating in this context owing to its biocompatibility, favorable pharmacokinetics, prolonged circulation and intrinsic capacity to associate with hydrophobic compounds and to accumulate in tumours [24–26]. The albumin-coated preparation enabled in vivo testing of the VS1–5-FU combination and was associated with improved tolerability, underscoring that formulation is a key determinant of translatability for this class of insoluble inhibitors.

Our data also reveal a selective functional interaction between STARD3 inhibition and fluoropyrimidine treatment. Among the tested CRC drugs, VS1 showed the most consistent synergistic effect with 5-FU across cell models, whereas combinations with oxaliplatin, irinotecan or regorafenib were less favorable. This selectivity suggests that STARD3 blockade does not act as a general chemosensitizer but instead interacts with vulnerabilities that are particularly relevant to fluoropyrimidine response. Although the precise mechanism remains to be fully established, several non-mutually exclusive explanations can be considered. By perturbing intracellular cholesterol trafficking, STARD3 inhibition may alter membrane organization and lipid raft composition, thereby affecting receptor signaling and transporter function [2,11,12,47,48], These changes could in turn influence intracellular drug retention or adaptive stress responses that contribute to 5-FU resistance. In parallel, disruption of cholesterol homeostasis may sensitize cells to mitochondrial dysfunction and apoptotic priming. In line with this, we observed that the combination selectively drives a significant increase in ROS production only in synergy-responsive cell lines. This suggests that the disruption of cholesterol homeostasis directly fuels oxidative stress and apoptotic priming, thereby acting as a key driver of the observed synergistic cytotoxicity [49,50]. Given the known impact of 5-FU on nucleotide metabolism, STARD3 blockade may also exacerbate metabolic stress and replication-associated damage [51–53]. These mechanistic links remain inferential at this stage and will require direct experimental validation, but they provide a plausible framework for the observed interaction.

Importantly, the combination of VS1-AF and 5-FU produced a stronger antitumor effect in vivo than either monotherapy alone, as evidenced by reduced tumor growth, prolonged doubling time and growth delay, increased intratumoral necrosis, and absence of aggravated systemic toxicity.

Our findings also have potential implications for biomarker-guided treatment. STARD3 has been linked to aggressive disease features and poor outcome in several tumor types [18–22] raising the possibility that tumors with elevated STARD3 expression or heightened cholesterol trafficking dependency may be particularly responsive to this strategy. In this respect, the present study provides a proof-of-concept for targeting cholesterol transport protein to improve chemotherapy efficacy. Future work should determine whether STARD3 levels, localization, or broader lipid-metabolic signatures can identify CRC subsets most likely to benefit from VS1-based combinations.

In summary, we demonstrate a carrier-free, albumin-coated nanocrystal formulation that converts the insoluble, first-in-class STARD3 inhibitor VS1 into an in vivo-active agent. The formulation increases apparent aqueous solubility more than 14-fold, provides sustained dissolution-controlled release, and preserves the crystallinity and molecular integrity of the drug. Delivered in this form, VS1 selectively potentiates 5-FU in 5-FU-sensitive CRC models and, combined with 5-FU, suppresses xenograft growth without aggravating systemic toxicity. By coupling a structurally defined inhibitor with a simple, scalable delivery platform, this work provides a formulation-enabled route to combination strategies against fluoropyrimidine-resistant CRC and a template for advancing other poorly soluble oncology probes toward in vivo evaluation.

## DISCLAIMER

During manuscript preparation, the authors used ChatGPT solely for language editing. The authors reviewed and edited all AI-assisted text and took full responsibility for the final content.

## ACKNOWLEDGMENTS

We thank the XRD2 beamline at Elettra-Sincrotrone Trieste for beamtime.

## DATA AVAILABILITY

The data supporting the findings of this study are available in the paper, supplementary information files, and can be obtained from the corresponding authors upon reasonable request. The accession number for the coordinates and structure factors for the VS1-bound form of StARD3LBD domain reported in this paper is PDB: 9RUE.

## AUTHOR CONTRIBUTIONS

I.C.: conceptualization, data analysis, review & editing; A.B.: experimental and data analysis and writing; G.S.: experimental and data analysis and writing; L.M.R.N.: experimental and data analysis and writing; G.P.: experimental and data analysis and writing; S.B.K.: experimental and data analysis; U.K.: experimental and data analysis and writing; M.DS.: experimental and data analysis; K.S.S.: experimental and data analysis; J.L.P: experimental and data analysis; R.H.: data collection; S.P: experimental and data analysis; J.B.: data analysis; M.A.: conceptualization; C.G.: review & editing; M.DS.: review & editing; S.O.: review & editing; M.C.: review & editing; T.T.: data analysis, review & editing; V.C.: review & editing; F.R: conceptualization, data analysis, review & editing. All authors reviewed and approved the final manuscript.

## FUNDING ACQUISITION

This research was funded to I.C. by the Ministry of Health, GR-2021-12374267. S.O. is supported by Fondazione AIRC per la Ricerca sul Cancro Investigator Grant IG20778. This research was also partially funded by the Slovenian Research and Innovation Agency (Program No P3-0003).

