## Supplementary Material for "Albumin-coated VS1 nanocrystals enable STARD3 inhibition and potentiate fluoropyrimidine therapy in colorectal cancer"

Prof. Flavio Rizzolio

Department of Molecular Sciences and Nanosystems

Ca' Foscari University Venice, Italy

+39 041 234 8934

Dr. Isabella Caligiuri

Pathology Unit

Centro di Riferimento Oncologico di Aviano (C.R.O.) IRCCS, Italy

+39 0434 659026

1 Pathology Unit, Centro di Riferimento Oncologico di Aviano (C.R.O.) IRCCS, 33081, Aviano, Italy

2 Department of Molecular Sciences and Nanosystems, Ca' Foscari University of Venice, 30172, Venice, Italy

3 Structural Biology Laboratory, Elettra-Sincrotrone Trieste, Trieste, Italy

4 Department of Pharmacy, University of Pisa, 56124, Pisa, Italy

5 Department of Experimental Oncology, Institute of Oncology Ljubljana, Zaloska cesta 5, SI-1000, Ljubljana, Slovenia

6 Department of Medical Oncology, Centro di Riferimento Oncologico di Aviano (C.R.O.) IRCCS, 33081, Aviano, Italy

7 Department of Medical, Surgical and Health Sciences, University of Trieste, 34127 Trieste, Italy

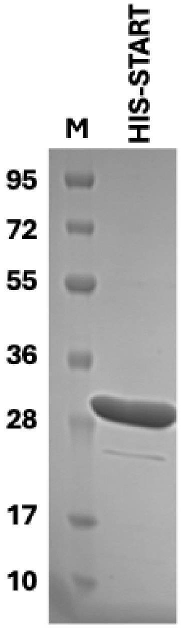

**Supplementary Figure S1. Purification of recombinant StARD3$_{\text{LBD}}$**.SDS-PAGE gel image of the recombinant START domain used for crystallographic studies. **M**, molecular weight marker.

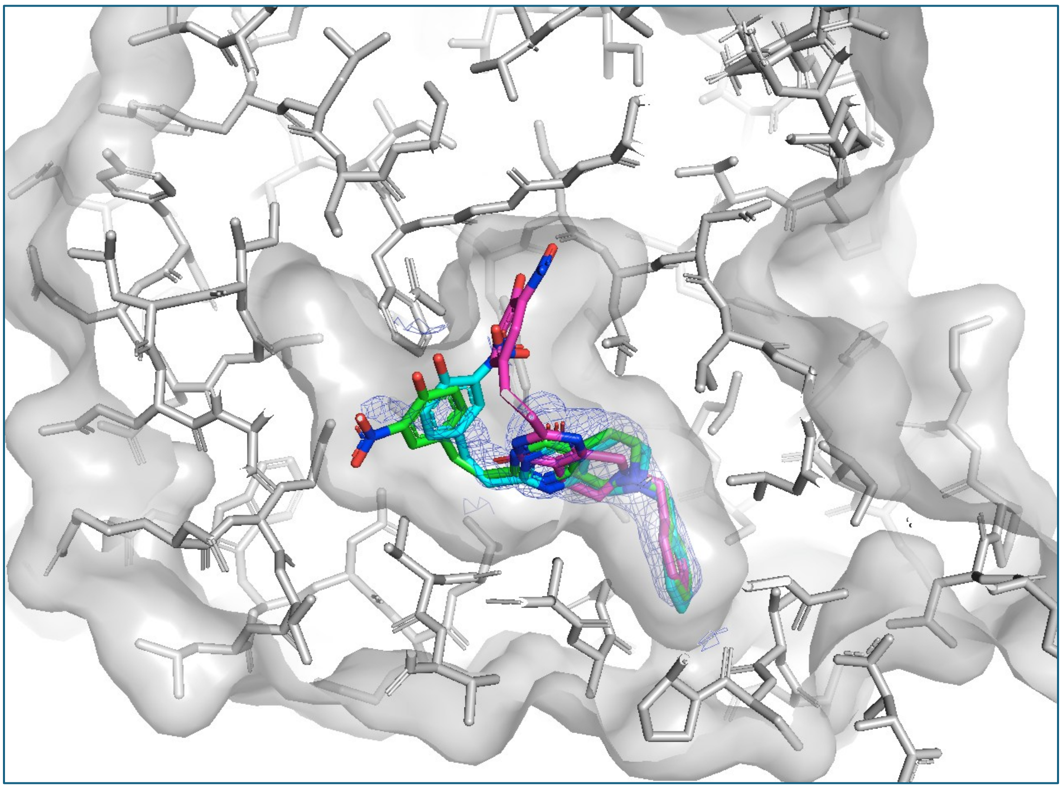

**Supplementary Fig. S2. Alternative conformations of VS1 in the STARD3 binding pocket**. Surface representation of the cavity and electron density around the inhibitor indicate flexibility of the nitrophenol moiety; conservative modeling was performed using a single conformation with partial occupancy.

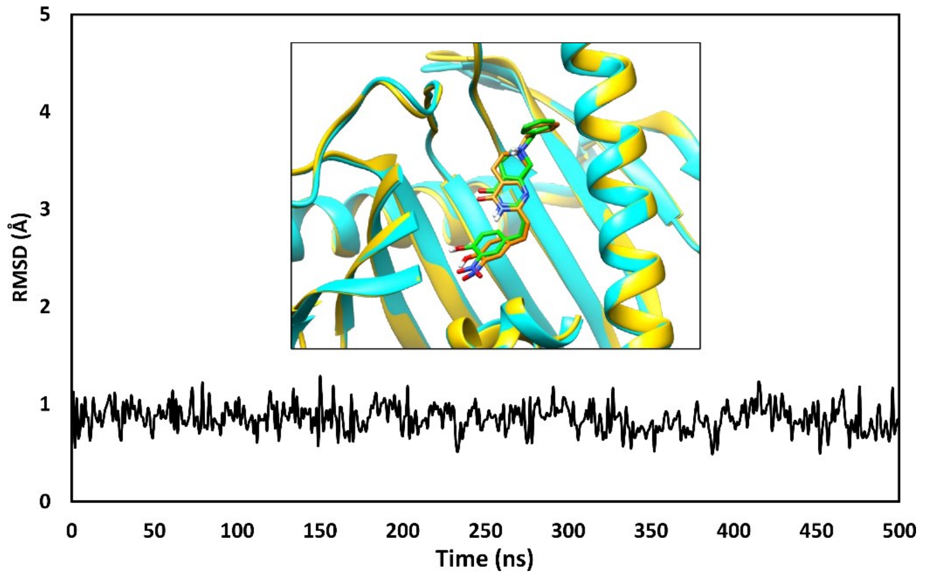

**Supplementary Fig. S3. Analysis of VS1 binding mode during classic MD**. The RMSD of the ligand disposition during the 500 ns of classic MD simulation is shown in black. The inner panel shows the superposition of the starting conformation of the ligand (green) into the protein binding site (cyan) with the average ligand conformation (orange) within STARD3 (gold) obtained from the last 200 ns of MD. For clarity, the α-helices and the β-sheets in the front have been removed to better show the inside of the cavity.

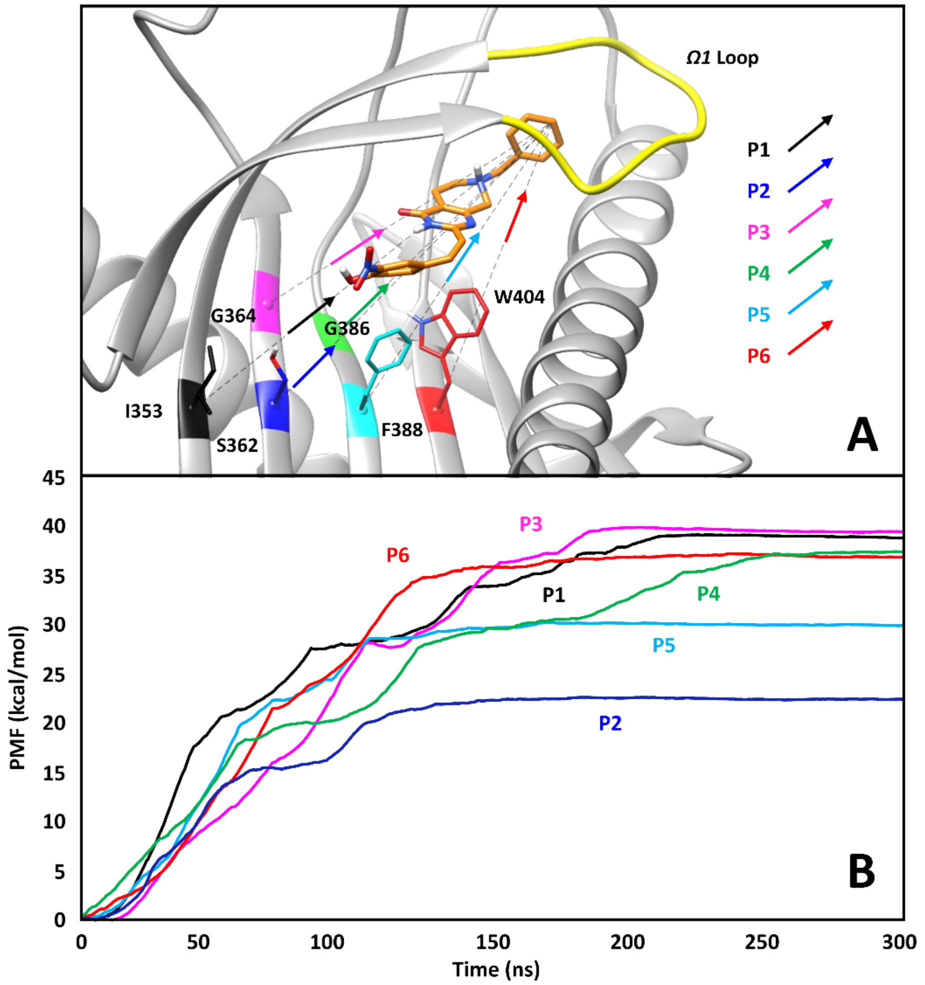

**Supplementary Fig. S4**. (**A**) **Starting frame of the SMD simulations**. The ligand is shown as orange sticks, while the protein is displayed as gray ribbons, with the exception of the Ω1 loop that is highlighted in yellow. The six different residues (I353, S362, G364, G386, F388, and W404), whose α carbons were used to guide the unbinding pathways (P1, P2, P3, P4, P5 and P6, respectively) are also displayed. For clarity, the α-helix and the β-sheet in the front have been removed to better show the inside of the cavity. The colored arrows show the initial pulling direction of each pathway. (**B**) Potential of mean force (PMF) curves corresponding to the free energy associated with the six different unbinding pathways.

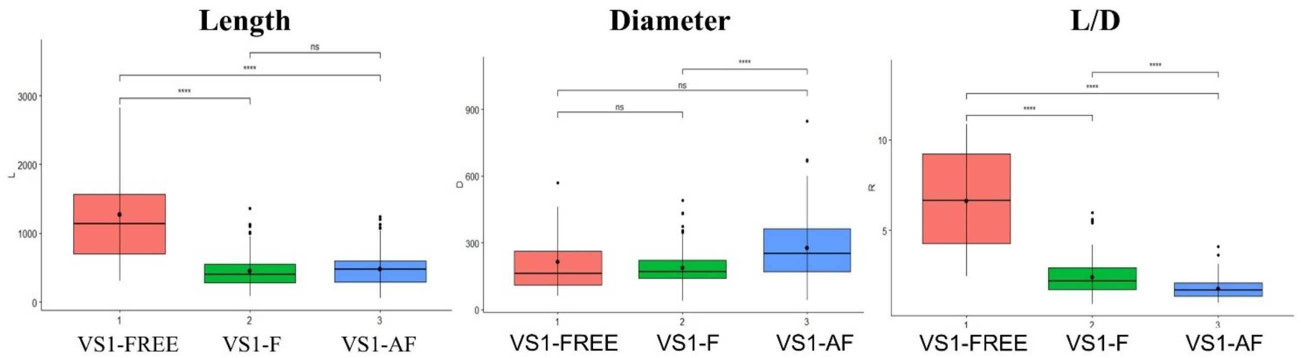

**Supplementary Fig. S5**: Boxplots to compare TEM measurement data for VS1 dimensions (Wilcoxon test p-value * <0.05, ** <0.01, ***<0.0001).

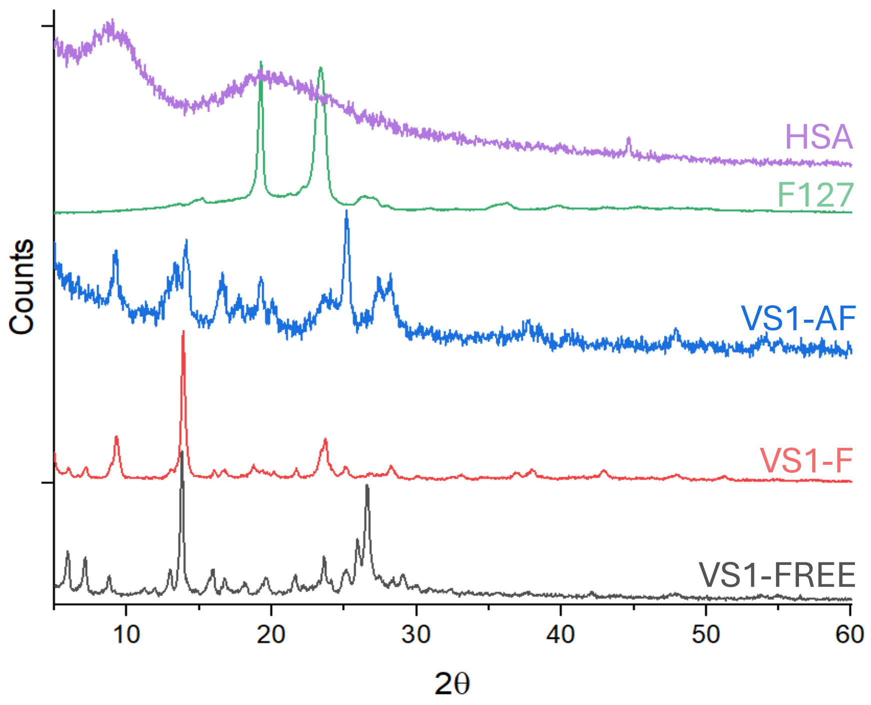

**Supplementary Fig. S6**: XRD of F127 powder, HSA powder, VS1 free i,e, drug powder, powder of VS1F and VS1-AF.

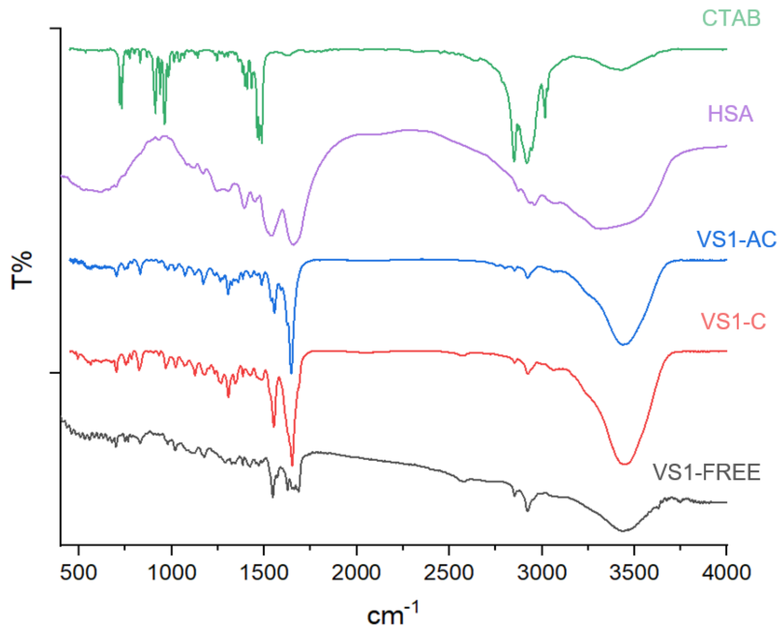

**Supplementary Fig. S7.** FTIR spectra of CTAB powder, HSA powder, VS1 drug powder, VS1-C powder and VS1-AC powder.

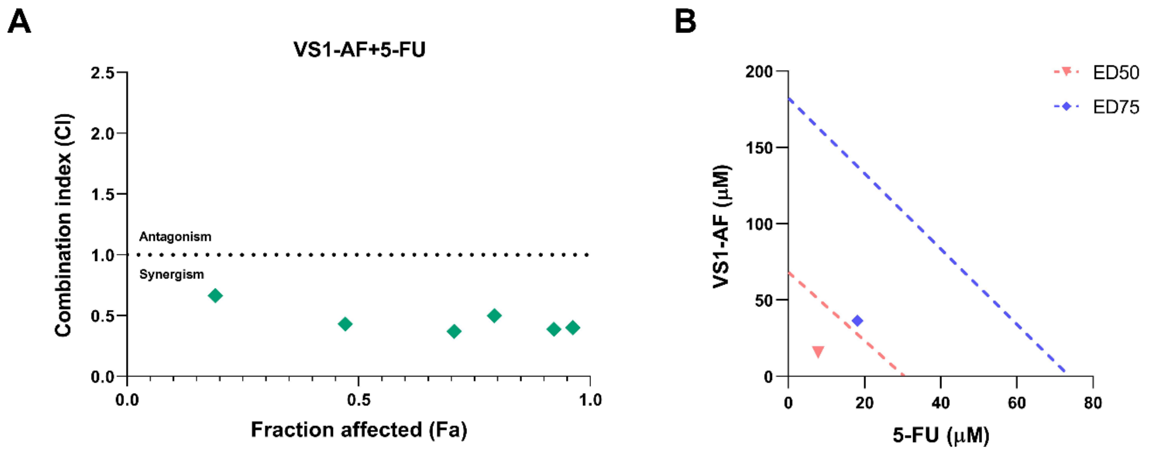

**Supplementary Fig. S8**: **Synergistic interaction between VS1-AF and 5-FU in HCT-116 cells**. (**A**) Fa-CI plot showing combination index (CI) values below 1 across all tested effect levels, indicating synergism. (**B**) Isobologram at ED50 and ED75, with data points falling below the line of additivity, confirming the synergistic interaction.

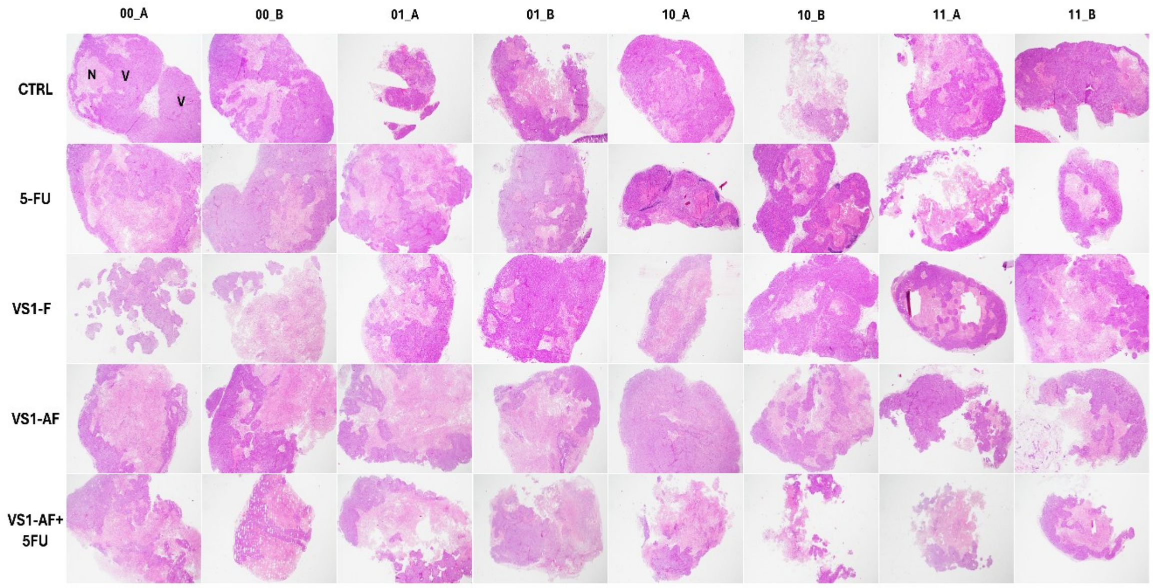

**Supplementary Fig. S9**: **Different proportion of tumor necrosis among the groups in cutaneous/subcutaneous tumor**. Vital (V) or necrotic (N) neoplastic tissue was indicated. Hematoxylin and Eosin staining. Original magnification × 2.

### Supplementary Tables

| Data collection | |
|---|---|
| Space group | P $3_1$ 2 1 |
| **Cell dimensions** | |
| *a*, *b*, *c* (Å) | 83.87 83.87 82.31 |
| α, β, ϒ | 90◦ 90◦ 120◦ |
| $R_{merge}$ | 0.064 |
| Resolution (Å) | 2.09 |
| I/σ(I) | 2.43 |
| Completeness (%) | 93.3 |
| Redundancy | 15.8 |
| **Refinement** | |
| No. reflections | 30380 |
| $R_{work}/R_{free}$ | 0.203/0.233 |
| No. atoms | |
| Protein | 1778 |
| Ligand/ion | 30/10 |
| Water | 156 |
| *B*-factors | 40.3 |
| Protein | 44.0 |
| **RMSD** | |
| Bond lengths (Å) | 0.008 |
| Bond angles (°) | 0.92 |

Values in parentheses are for the highest resolution bin.

**Supplementary Table S1. STARD3 X-ray data collection, processing, and refinement statistics.**

| VS1-free | VS1-F | | Molecular group |
|---|---|---|---|
| Positions ($cm^{-1}$ ) | Positions ($cm^{-1}$) | Δ ($cm^{-1}$) | |
| 1545 | 1552 | -7 | ν $NO_2$ |
| 1571 | X | | ν C=C |
| 1625 | 1651 | different multiplicity, intensity | ν CO |
| 1649-1686 | | | ν C=C + ν C=N + δ $H_2O$ |
| 2851 | 2849 | 2 | ν CH aliphatic |
| 2918 | 2920 | 2 | ν CH |
| 3446 | 3434 | 12 | ν OH + ν NH |

**Supplementary Table S2. FTIR peak assignments for VS1-free and VS1-F.**

| Sample | EE % |
|---|---|
| VS1-C | 93 ± 9 |
| VS1-AC | 60 ± 20 |

**Supplementary Table S3. Encapsulation efficiency and albumin content of CTAB-based formulations.**

| VS1-free | VS1-C | | Molecular group |
|---|---|---|---|
| Positions ($cm^{-1}$) | Positions ($cm^{-1}$) | Δ ($cm^{-1}$) | |
| 1545 | 1552 (with shoulder) | (with shoulder) | ν $NO_2$ |
| 1571 | x | | ν C=C |
| 1625 | 1626 | Shoulder, different intensity | ν CO |
| 1649-1686 | 1651 | different multiplicity, intensity | ν C=C + ν C=N + δ $H_2O$ |
| 2851 | 2849 | 3 | ν CH aliphatic |
| 2918 | 2919 | 1 | ν CH |
| 3446 | 3447 | 1 | ν OH + ν NH |

**Supplementary Table S4. FTIR peak assignments for VS1-free and VS1-C.**

| Drug combination (A + B) | 1 | 2 | 3 | 4 | 5 | 6 | 7 |
|---|---|---|---|---|---|---|---|
| VS1 - Regorafenib | 200 - 10 | 100 - 5 | 50 - 2.5 | 25 - 1.12 | 12.5 - 0.62 | 6.25 - 0.31 | 3.12 - 0.15 |
| VS1 - 5-Fluorouracil | 200 - 100 | 100 - 50 | 50 - 25 | 25 - 12.5 | 12.5 - 6.25 | 6.25 - 3.12 | 3.12 - 1.56 |
| VS1 - Oxaliplatin | 200 - 5 | 100 - 2.5 | 50 - 1.25 | 25 - 0.62 | 12.5 - 0.31 | 6.25 - 0.15 | 3.12 - 0.08 |
| VS1 - Irinotecan | 200 - 5 | 100 - 2.5 | 50 - 1.25 | 25 - 0.62 | 12.5 - 0.31 | 6.25 - 0.15 | 3.12 - 0.08 |

**Supplementary Table S5. Concentrations (μM) of drug combinations used for synergy evaluation.**

| Cell line | VS1 + 5-FU (μM) |
|---|---|
| HCT-116 | 50 – 4 |
| COLO-201 | 25 – 16 |

**Supplementary Table S6. Concentrations of VS1 and 5-FU used for ROS evaluation.**
